# Monitoring photogenic ecological phenomena: Social network site images reveal spatiotemporal phases of Japanese cherry blooms

**DOI:** 10.1101/2021.09.13.460016

**Authors:** Moataz Medhat ElQadi, Adrian G Dyer, Carolyn Vlasveld, Alan Dorin

**Affiliations:** Faculty of Information Technology, Monash University, Clayton, VIC 3800, Australia; School of Media and Communications, RMIT University, Melbourne, VIC 3000, Australia; Faculty of Science, Monash University, Clayton, VIC 3800, Australia

**Keywords:** social network site data, phenology, iEcology, incidental citizen science, spatiotemporal analysis

## Abstract

Some ecological phenomena are visually engaging and widely celebrated. Consequently, these have the potential to generate large footprints in the online and social media image records which may be valuable for ecological research. Cherry tree blooms are one such event, especially in Japan where they are a cultural symbol (Sakura, 桜). For centuries, the Japanese have celebrated Hanami (flower viewing) and the historical data record of the festival allows for phenological studies over this period, one application of which is climate reconstruction. Here we analyse Flickr social network site data in an analogous way to reveal the cherry blossoms’ seasonal sweep from southern to northern Japan over a twelve-week period.

Our method analyses data filtered using geographical constraints, multi-stage text-tag classification, and machine vision, to assess image content for relevance to our research question and use it to estimate historic cherry bloom times. We validated our estimated bloom times against official data, demonstrating the accuracy of the approach. We also investigated an out of season Autumn blooming that has gained worldwide media attention. Despite the complexity of human photographic and social media activity and the relatively small scale of this event, our method can reveal that this bloom has in fact been occurring over a decade.

The approach we propose in our case study enables quick and effective monitoring of the photogenic spatiotemporal aspects of our rapidly changing world. It has the potential to be applied broadly to many ecological phenomena of widespread interest.

## 1 Introduction

Climate change is disrupting many ecological phenomena, threatening insect pollinators vital to our food supply [1], generating conditions that increase the likelihood of wildfires [2], raising sea levels [3], and disrupting species’ viable ranges [4]. The rate, extent and diversity of these changes pose a challenge to traditional ecological observation and data collection methods. However, new sources of data are becoming available and increasingly important in this context. In this paper, we explore the application of social network site (SNS) data to monitor spatiotemporal ecological phenomena with high public profiles or a visibly engaging photographic presence. Our case study is the blooming of Japanese cherry trees.

Today, humanity documents its activities digitally in the form of online scientific publications (e.g. elsevier.com), entertainment content (e.g. Netflix.com, Youtube.com), financial data (e.g. NYSE.com), posts of personal interests (e.g. Twitter.com, Facebook.com), and physical activity (e.g. Strava.com). The unprecedented quantity and variety of data online has drawn researchers’ attention for applications ranging from public opinion and sentiment analysis [5] and election forecasting [6], to natural disaster monitoring [7], remote sensing [8], epidemiology [9], and ecology [10, 11].

Geo-tagged visual media in particular, as a form of volunteered geographic information, has seen strong interest in scientific research. This data may include photographic evidence of events in remote areas that would otherwise be impractical to survey [10–14], but it may also contain observations of popular events, in which case the sheer abundance of the data is potentially of benefit. Daume [12] however, has noted that although Twitter, the source of data in his study of invasive species, is potentially a rich source of observations, there are technical challenges involved in using SNS data; careful validation of results is required. There has also been concern raised about the implications for personal privacy of the availability of mass data sets being used in research, especially when the subject of the research relates to the personal attributes of people uploading images of themselves and their friends [15].

Of specific relevance to the work presented in this paper, if carefully managed, volunteered information gleaned from SNS may provide a valuable source of data to monitor and understand the dynamics of ecosystems [16, 17]. In effect, every person posting to a social network site might be what we propose to call an “incidental citizen scientist” of value to ecological projects. Previous applications in the domain include a project where manually analysed tweets from a sample spanning three years were used to detect invasive species [12]. Purkart and Depa detected invasive species in new sites in Slovakia, Czechia and Austria using crowd-sourced information on Facebook [18]. In addition, Becken and Stantic monitored sentiments on the Great Barrier Reef in Australia using data contained in tweets [19].

The primarily socially-governed (rather than scientifically-governed) data-collection behaviour of incidental citizen scientists requires researchers to carefully assess the quality of their data [12]. In addition, researchers must be mindful that the data may be inadequate or unsuitable for answering some types of pertinent question. For example, the effect human activity is having on the climate, and the impact of this on ecosystems, is of major concern to ecologists [1], but it is not immediately apparent how interest in understanding these changes is reflected in the activities of SNS users (e.g., see [20]). How then can SNS data address such specific concerns?

There are a number of ecological climate-influenced phenomena that are highly visible and of ongoing public interest. This interest can potentially serve ecologists well since key climate indicators may be recorded in the SNS data record. Some of these phenomena are the subject of tourism-related activities, others are the “poster children” of television nature documentaries. Examples include the “Penguin Parade” of little penguins in southern Australia and their diurnal return to the beach after feeding [21], the mass migration of the monarch butterfly [22], the tourism surrounding autumn leaf viewing in the USA, Canada and Japan (“leaf peeping”, or in Japan, “momijigari”) [23], and, the spring bloom of cherry blossoms in Japan (“hanami”) [23]. All these phenomena are interesting for their own sake, visually impactful, but also, they are potentially helpful events by which to monitor biological and ecological events. Their general popularity generates a strong footprint online consisting of photographic records and social media posts, many of which are geo-tagged. In this paper, we examine the phenology of cherry trees, a traditional cultural hallmark of spring in Japan, through the lens of SNS Flickr users. The potential of SNS data for analysing cherry bloom-related tourist activity in Japan has previously been demonstrated by Endo and Hirota [24], who used geo-tagged tweets and data interpolation to provide real-time information on the best times and locations to observe blossoms.

Although long-term ecological empirical data is difficult to collect, if such data is available, the timing of plant phenological events can inform studies of environmental change since climate variation affects plant phenology [25, 26]. Thanks to their cultural significance, records of blooming activities of Japan’s cherry trees have been preserved for centuries in diaries and chronicles making them a valuable indicator of climatic variation [27]. This long, written, historical bloom record was used to reconstruct seasonal temperature variation over several centuries in Tokyo [28, 29] and Kyoto [27].

Here we hypothesise that SNS data, like that previously used for short-term tourist activity monitoring noted above, can be used to track long-term national cherry blooms, as was previously conducted using traditional chronicle data. In addition to studying the Japanese national cherry bloom, as in the previous studies we shall analyse the historical blooms of trees in the Hanami hotspots of Tokyo and Kyoto. We propose to do this using the flowering record encapsulated in Flickr photos 2008-2018. To benchmark our approach, we compare our estimates with the full bloom dates published by the Japan National Tourism Organisation (JNTO) sourced from the Japan Meteorological Corporation (JMC). After establishing the fidelity of our filtered and processed SNS data to historical data, we investigate an apparent anomaly in the data that suggests cherry blossoms have not only been blooming as expected in spring over the past decade, but also surprisingly in autumn.

## 2 Materials and Methods

Our approach to collecting cherry blossom flowering event data can be divided into two steps 1) data extraction from the social network site, entailing several distinct sequential stages, and then 2) filtration of data for relevance using increasingly finer resolutions. We report in this section on the results of each intermediate stage to simplify the explanation of our filtration process. Results obtained from analysis of the final filtered data set are provided in section 3.

### 2.1 Initial SNS site search

Although our methodology is agnostic to the data source, as noted in earlier chapters, we chose Flickr since its Application Programming Interface (API) allows for simultaneous temporal, spatial, and textual filters, simplifying our initial data collection process. We collected geo-tagged photos from Flickr by searching for the keyword “cherry blossom” using the python API client [30]. Our search was confined to photos taken in Japan, geo-located within a bounding box (lat. 31.186°– 46.178°, and long. 129.173°– 145.859°), in the time span 1 January 2008 to 31 December 2018, an extended period prior to the recent Covid-19 induced declines in global tourism [31]. This data was then filtered by masking against the geographic boundary of Japan obtained from gadm.org. A total of 80,915 photos remained (Fig 1).

**Fig 1.**
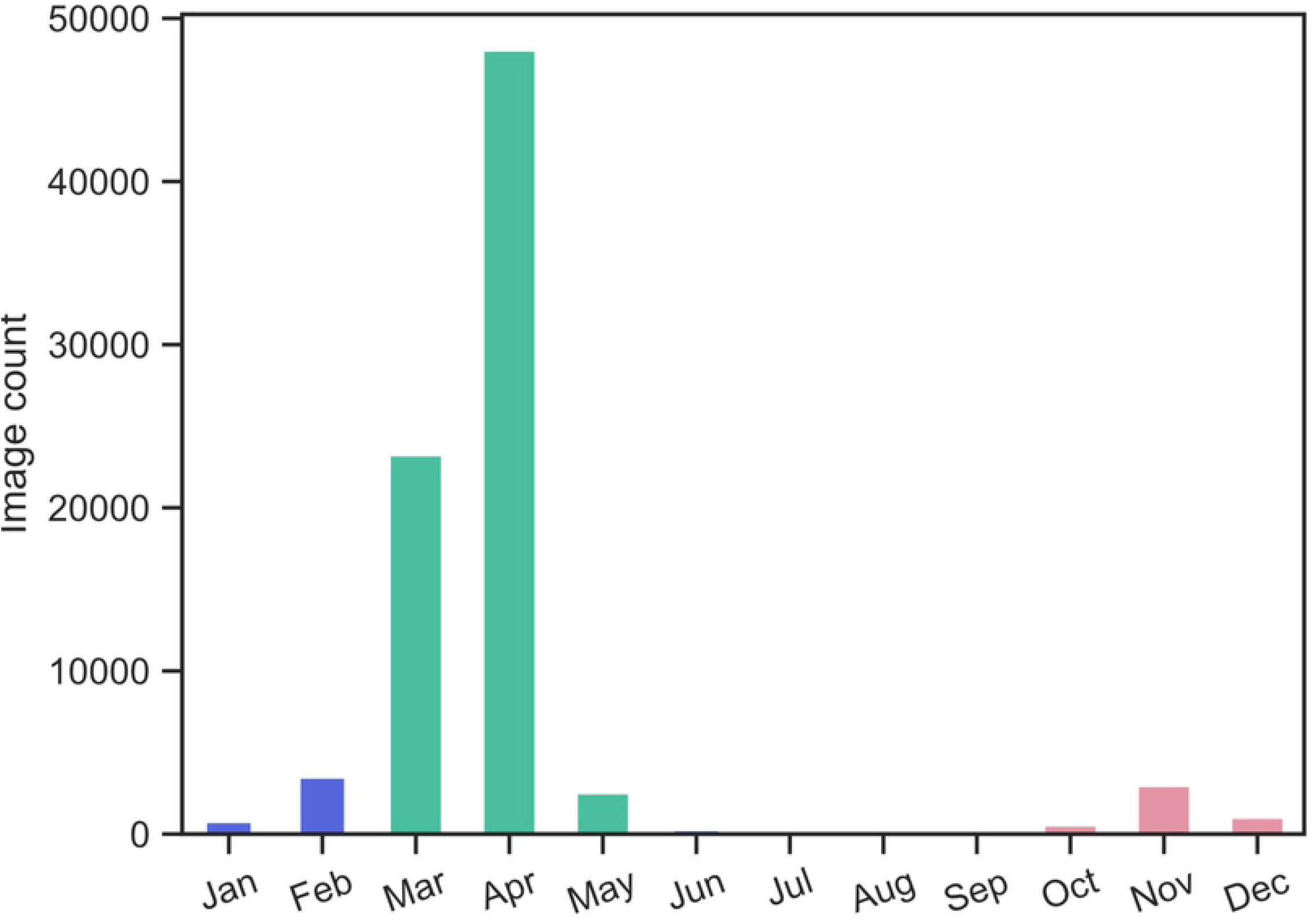
Total count of search results of “cherry blossom” photos from Flickr in Japan. Data range from 1 January 2008 to 31 December 2018 inclusive are shown. Data are concentrated in spring, but show an interesting small spike in autumn, peaking in November.

### 2.2 Filtration by computer vision generated tags

Although the Flickr photos were constrained using the search term “cherry blossom”, SNS content is frequently falsely (or only loosely) associated with search terms [10, 32]. Hence, we subjected all photos to a computer vision API that generates descriptive text tags for each photo based on its visual content, as a means to automatically double-check the relevance of individual data points. We used Google’s computer vision API (cloud.google.com/vision/) for this purpose. This API uses pre-trained machine learning models to assign labels to images based on predefined categories.

As anticipated, the computer vision API returned the text tag “cherry blossom” for most photos. Human analysis of the other returned tags revealed that most were conceptually related to cherry blossoms, except perhaps for the tags “autumn” and “maple tree”, that appeared in the last months of the year associated with some photographs (Fig 2). We might sensibly expect these tags to be associated by the computer vision algorithm with autumn leaves and Japanese Maples. A visual inspection of images with these tags confirmed this, from which we learnt that: (i) the computer vision API was correct in its assignment of the tag; and (ii) the data originally downloaded from Flickr using the search term “cherry blossom” contained images irrelevant to our goal. To refine the data set, we therefore chose to automatically maintain only photos with the tag “cherry blossom” and without tags “autumn” or “maple tree”.

**Fig 2.**
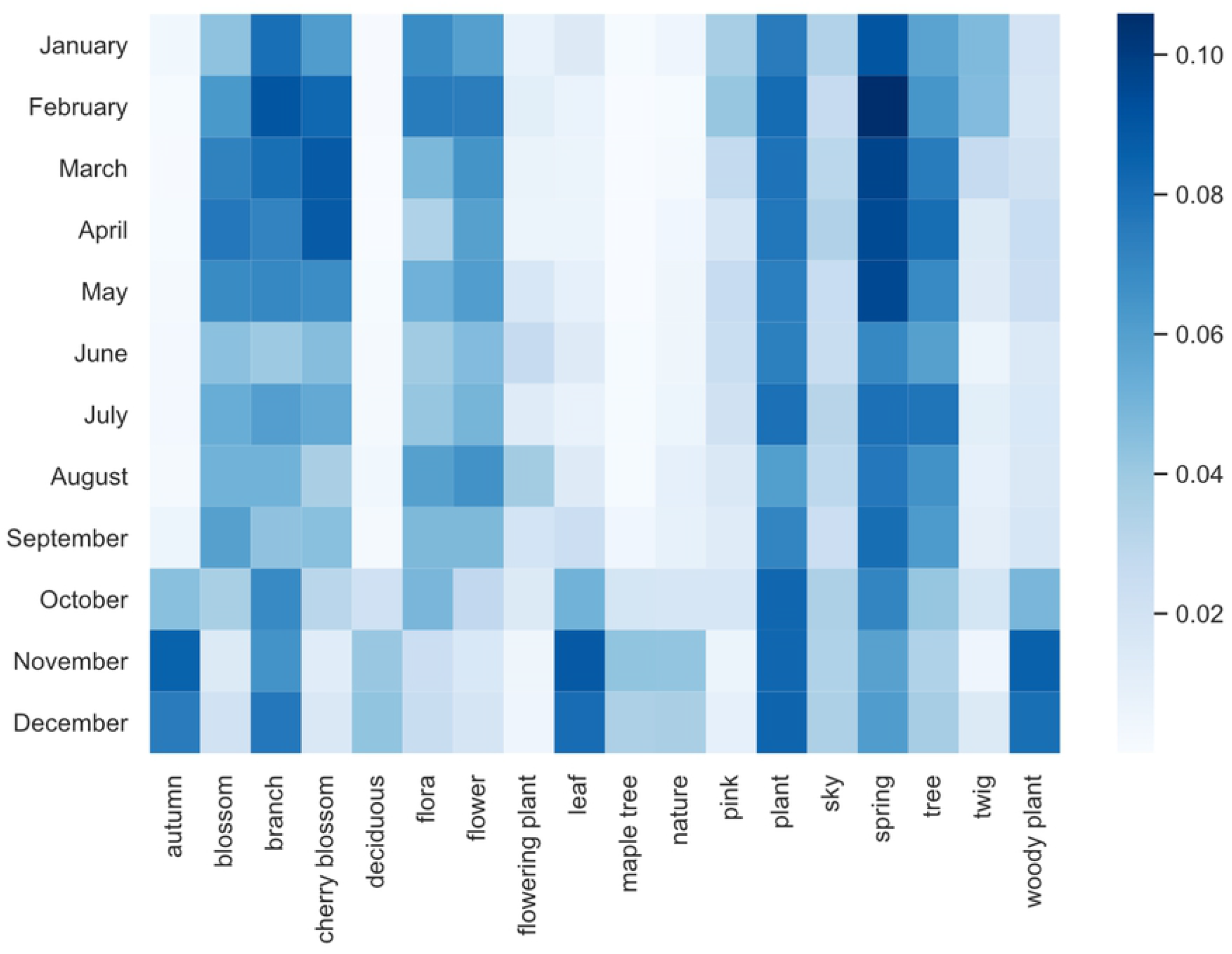
Normalised frequency of computer vision tags assigned to photos returned by Flickr search “cherry blossom”. Data are Japan-wide for the period January 2008 to December 2018 inclusive. Images within tags are not mutually exclusive; an image may be labelled simultaneously by several tags. To generate this plot, the top-10 most frequent text tags returned for images in each month were collected into a set of 18 tags. These appear alphabetically along the x-axis. The square marked for each month against a tag is coloured according to the relative frequency of its prevalence in that month. Note that “autumn” is a frequent tag in October, November and December. These photos were found by manual checking to contain autumn leaves rather than blossoms.

### 2.3 A closer look at Tokyo

After the initial data collection (section 2.1) and filtration (section 2.2) of photos sourced from across Japan (Fig 1), we used the geographic data associated with each image in the set to focus specifically on the region of Tokyo (district boundaries: gadm.org). This restricted our analysis to a built environment, which, along with climate generally, is important in considering tree phenology since temperature differences between rural areas and built environments can be several degrees due to the heat-island effect [33]. Tokyo represents a single climate zone and region of touristic interest, climate generally being a key determinant of cherry tree phenology [34], and tourism being a key determinant of photographic and social media activity (see Discussion, section 4).

To confirm the effectiveness of our text-tag based filter, a botanist (author C.V.) visually determined whether photos remaining in the data set depicted cherry blossoms. To achieve this, they checked in sequence whether each photo: 1) included a plant/tree; 2) if the plant/tree held flowers 3) if the flowers were identifiable as cherry blossoms (as distinct, for instance, from plum blossoms). We dismissed any photographs that failed any of the steps, or those which were of such poor quality or low resolution, that their content could not be accurately identified. These were omitted from the dataset as part of our method checking, and to ensure what remained was indeed human-validated evidence of cherry blossom activity. The sequence of image filtering and checking activities is documented in Results.

The visual similarity between cherry blossoms and other trees in the same genus, and also the inevitable lack of familiarity that international tourists may have with the cherry blossoms, can both possibly account for the misidentification of many cherry blossoms as plum blossoms. In each photograph, we distinguished cherry blossoms (*Prunus* subgenus *Cerasus*; Kato et al. 2014) from plum blossoms (*P. mume*) by their retuse petal apices, pedicellate flowers, horizontal bark lenticels and green leaves [35, 36]. Plants were identified as plum blossoms (and therefore not validated as cherry blossoms) by their rounded petal apices, round buds and nearly sessile flowers [36, 37]. A small number of other plants also misidentified as cherry blossoms were easily distinguishable as being from genera other than *Prunus* (e.g. *Wisteria*).

### 2.4 Full bloom date estimation

To estimate full bloom dates, we generated a one-day resolution time series for all photos within the final image data set. To this we applied a triangular rolling average of 7-day width (centroid ± 3 days, 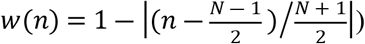 to smooth human photographic activity fluctuations occurring for Japanese local tourist activity between weekdays (typical working periods) and weekends (typical recreational periods). An example is provided (Fig **3**). The crests of photographic activity on the resulting graphs were identified as the peak bloom times according to our method and appear in our Results.

**Fig 3.**
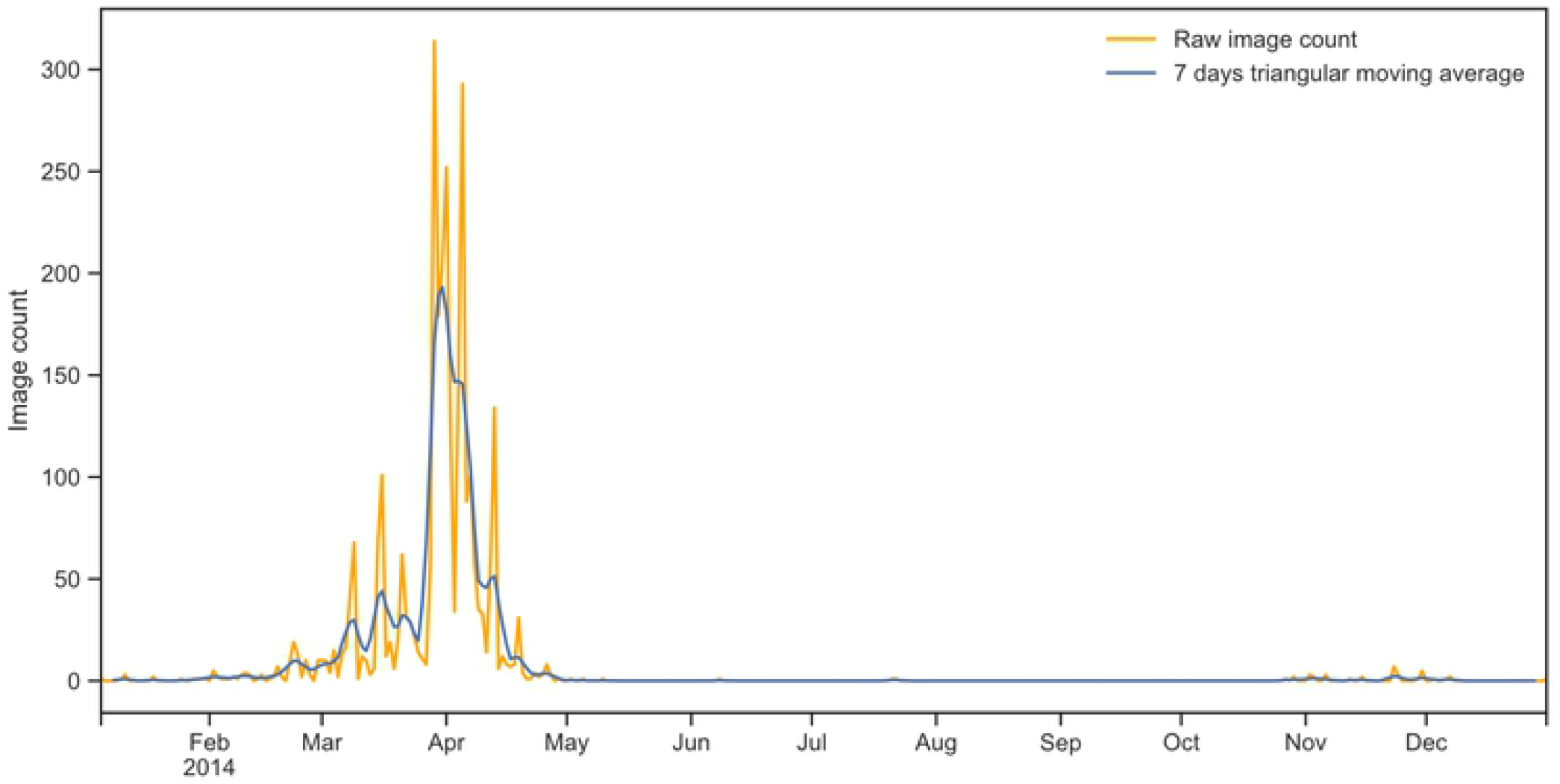
Estimating cherry full bloom date in Tokyo 2014 from Flickr image count. Raw image count is smoothed using a 7-day triangular moving window giving, for this example year, a peak on 29 March 2014.

### 2.5 Kernel density maps

To show the spatial density distribution of cherry blossom photos, we created kernel density maps using ArcGIS 10.5 [38] Kernel Density geoprocessing tool based on a quartic kernel function [39].

## 3 Results

Our results are presented for Japan overall (section 3.1), for Tokyo (sections 3.2 – 3.4), and for Kyoto (section 3.5).

### 3.1 National cherry bloom spatiotemporal data visualisation

The filtered dataset of geo-tagged cherry blossom photos from spring months extracted across the whole of Japan were overlayed on a geographical map of the country (Fig 4) using the method described in section 2.5. Each panel shows the locations of photos taken within a 2-week interval. The map shows a clear pattern that starts with photographic activity data of blooming in the warmer southern population centres and advances northward.

**Fig 4.**
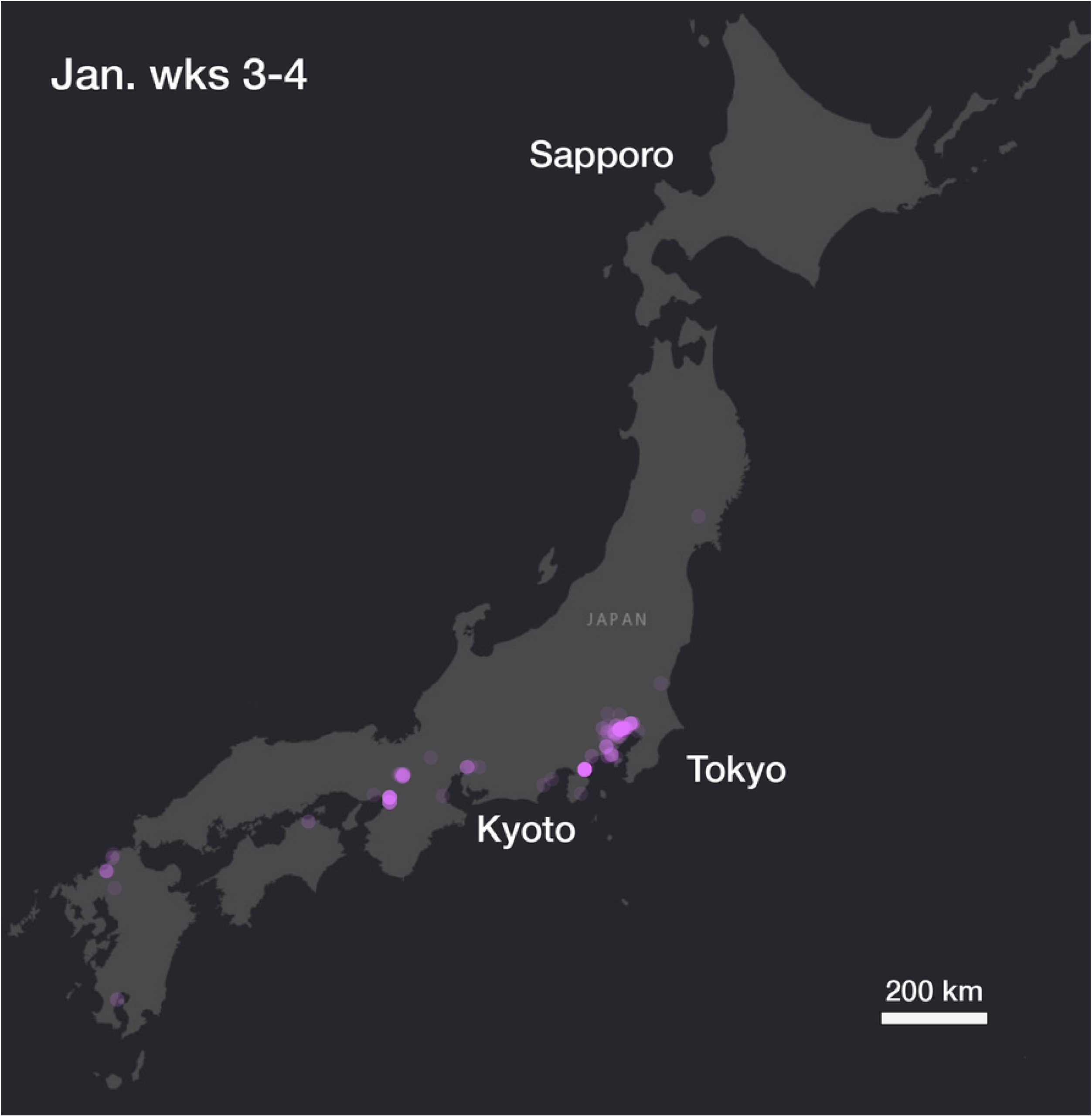
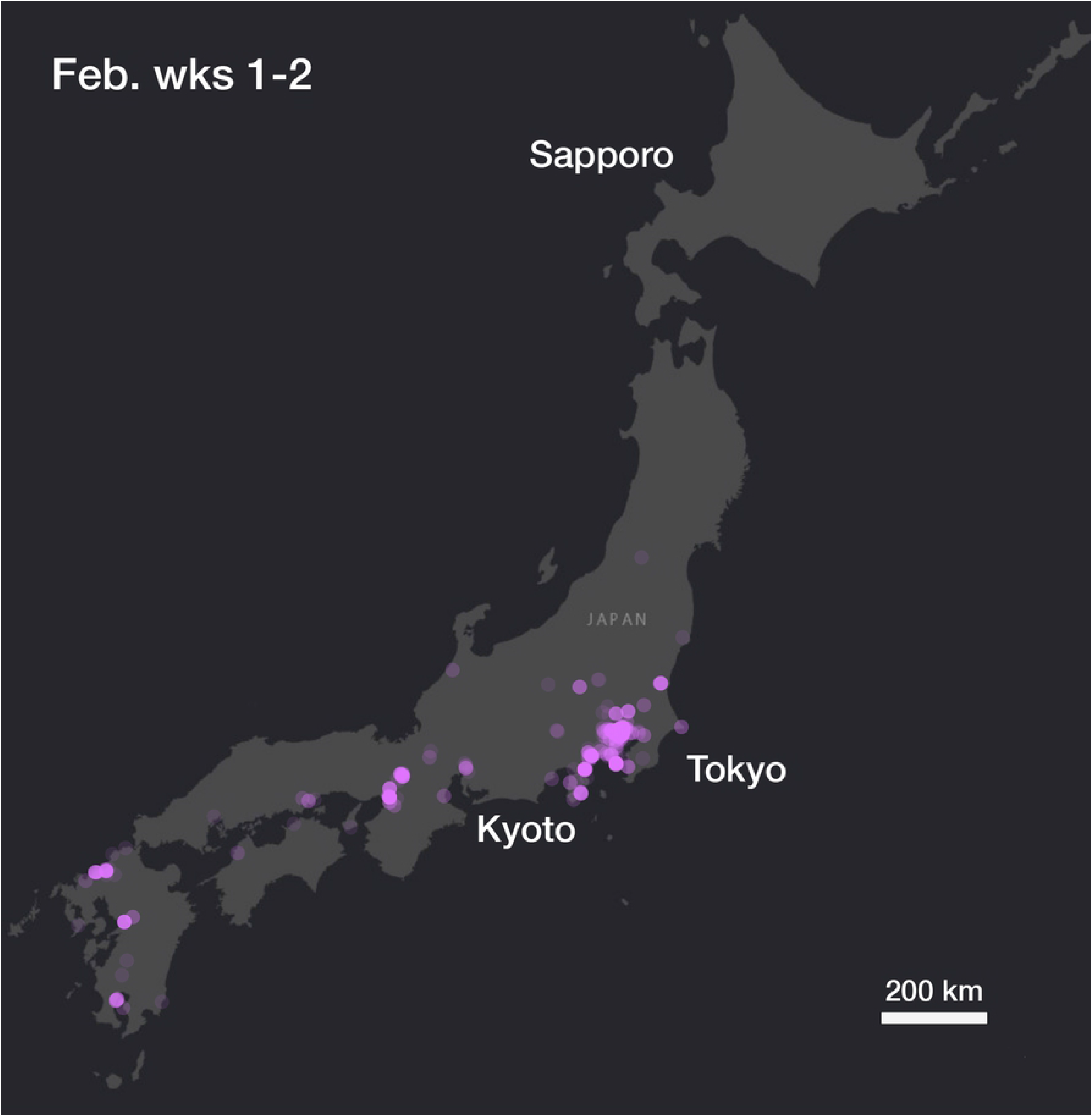
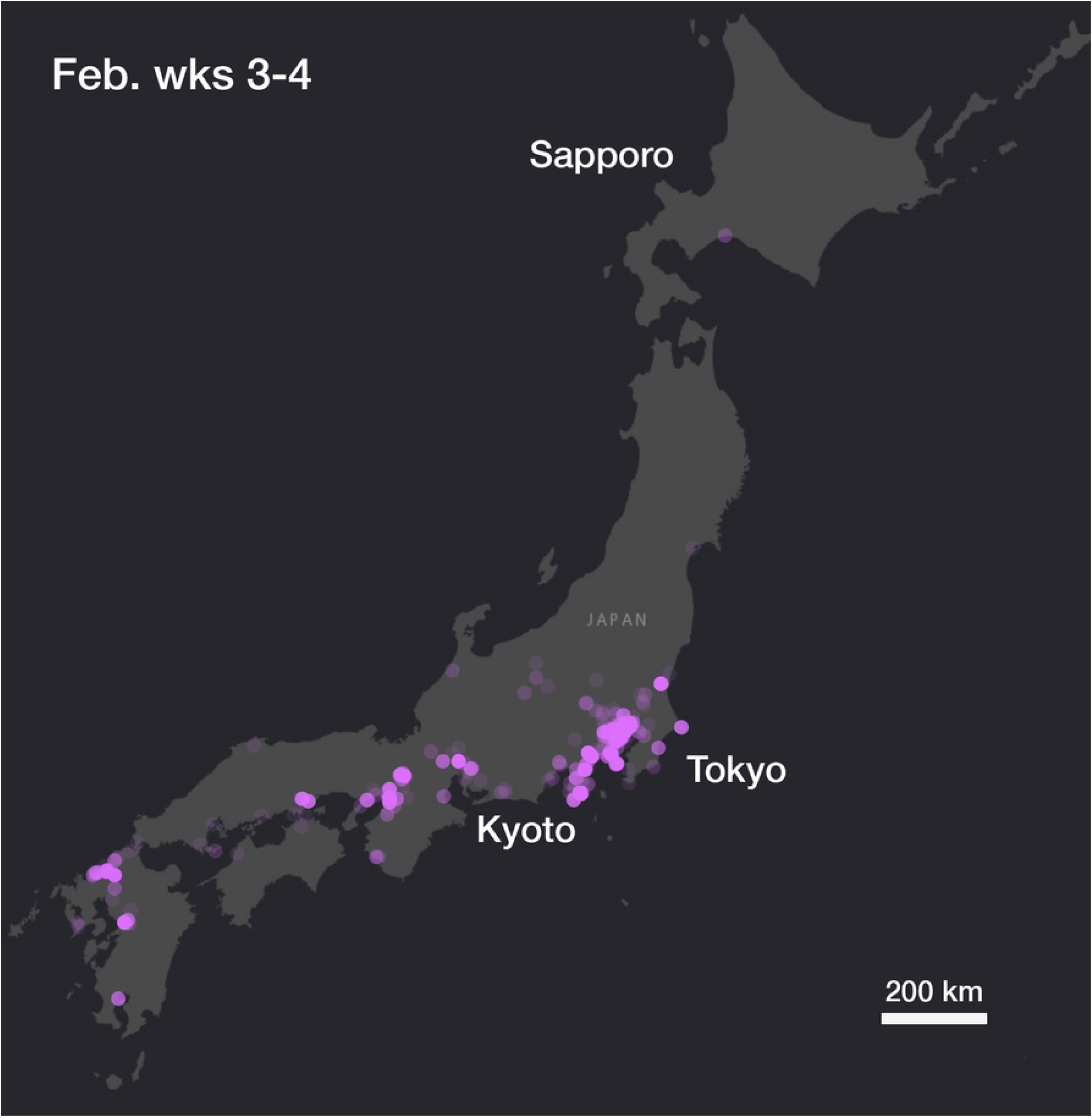
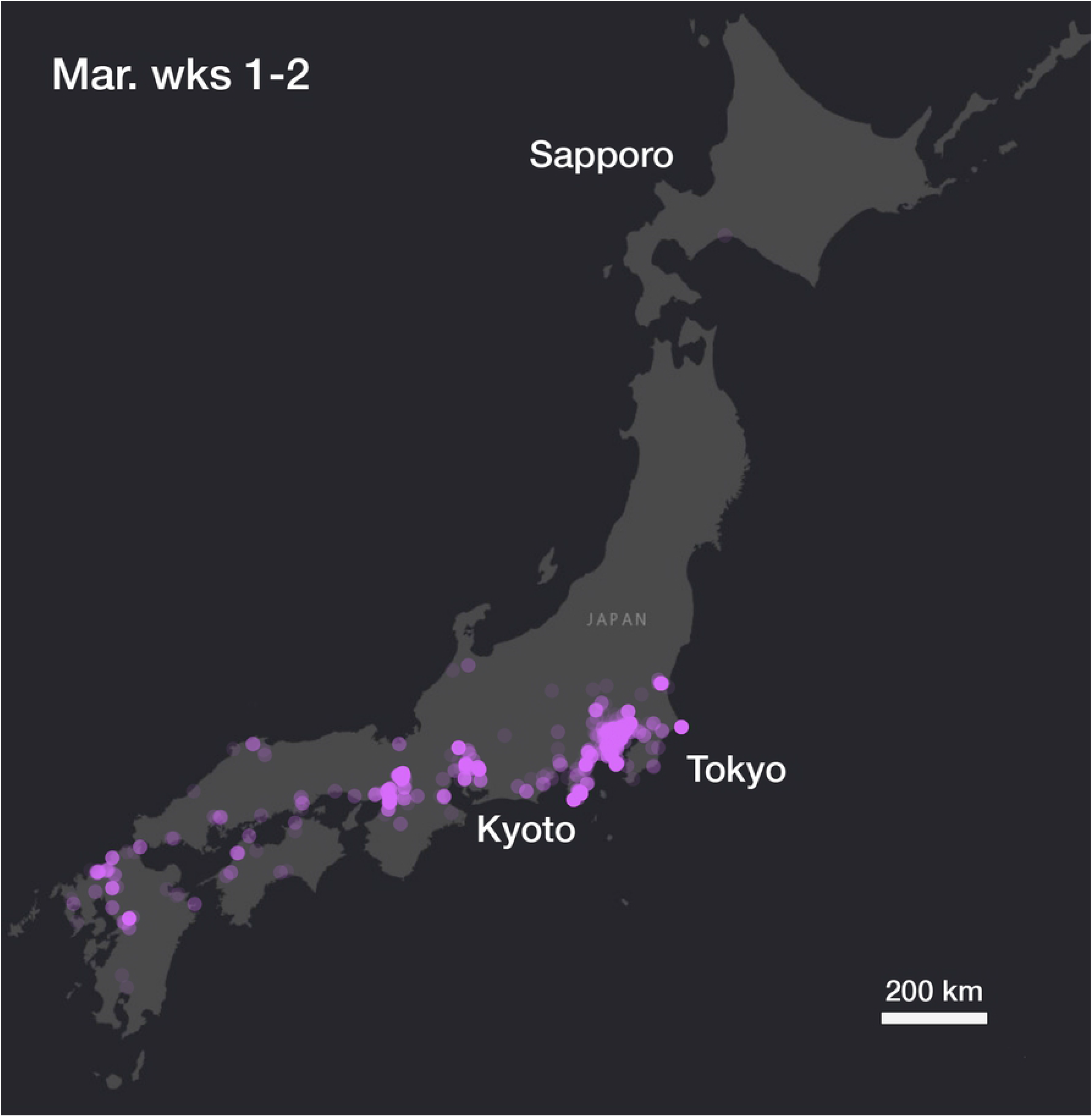
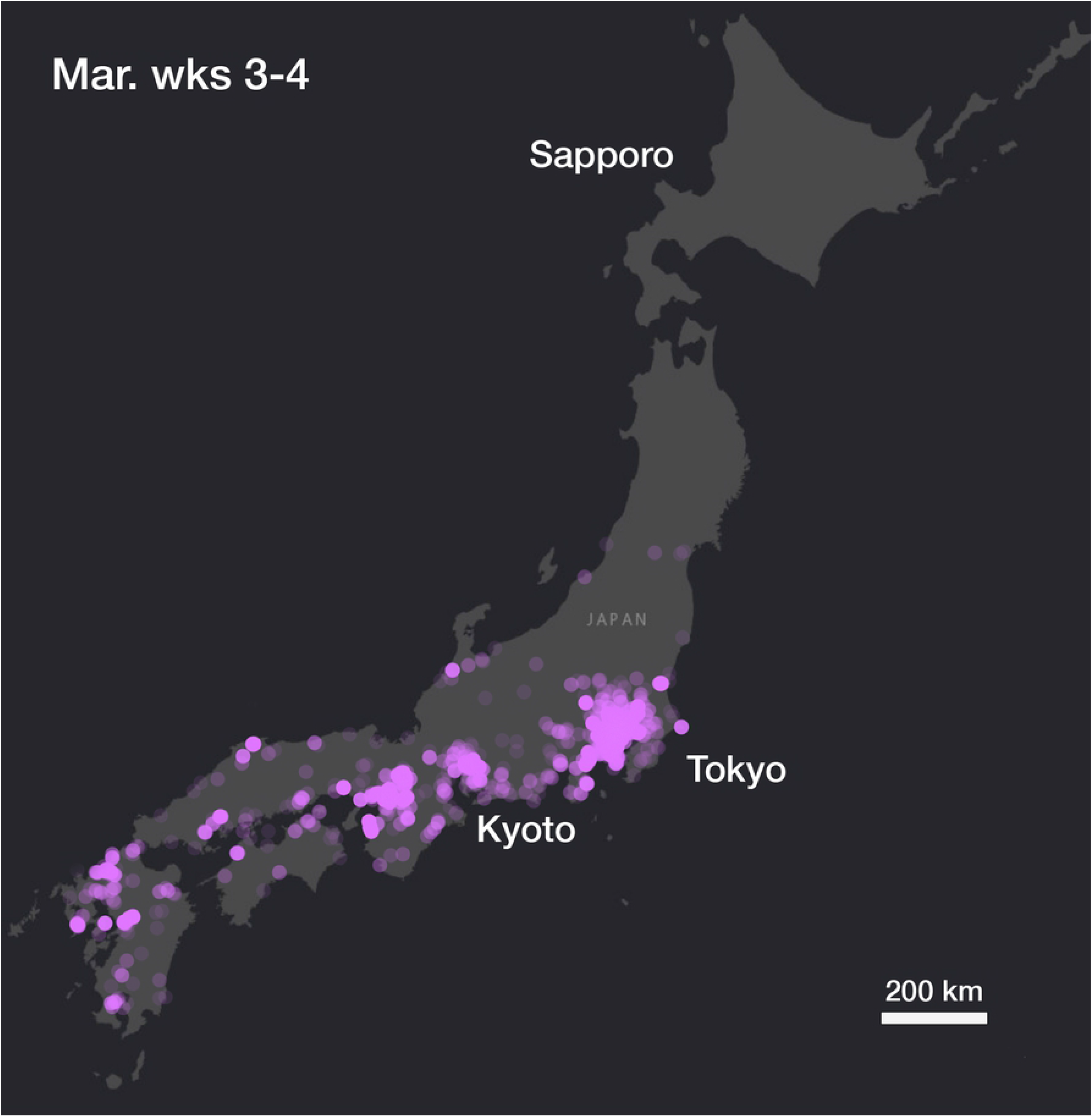
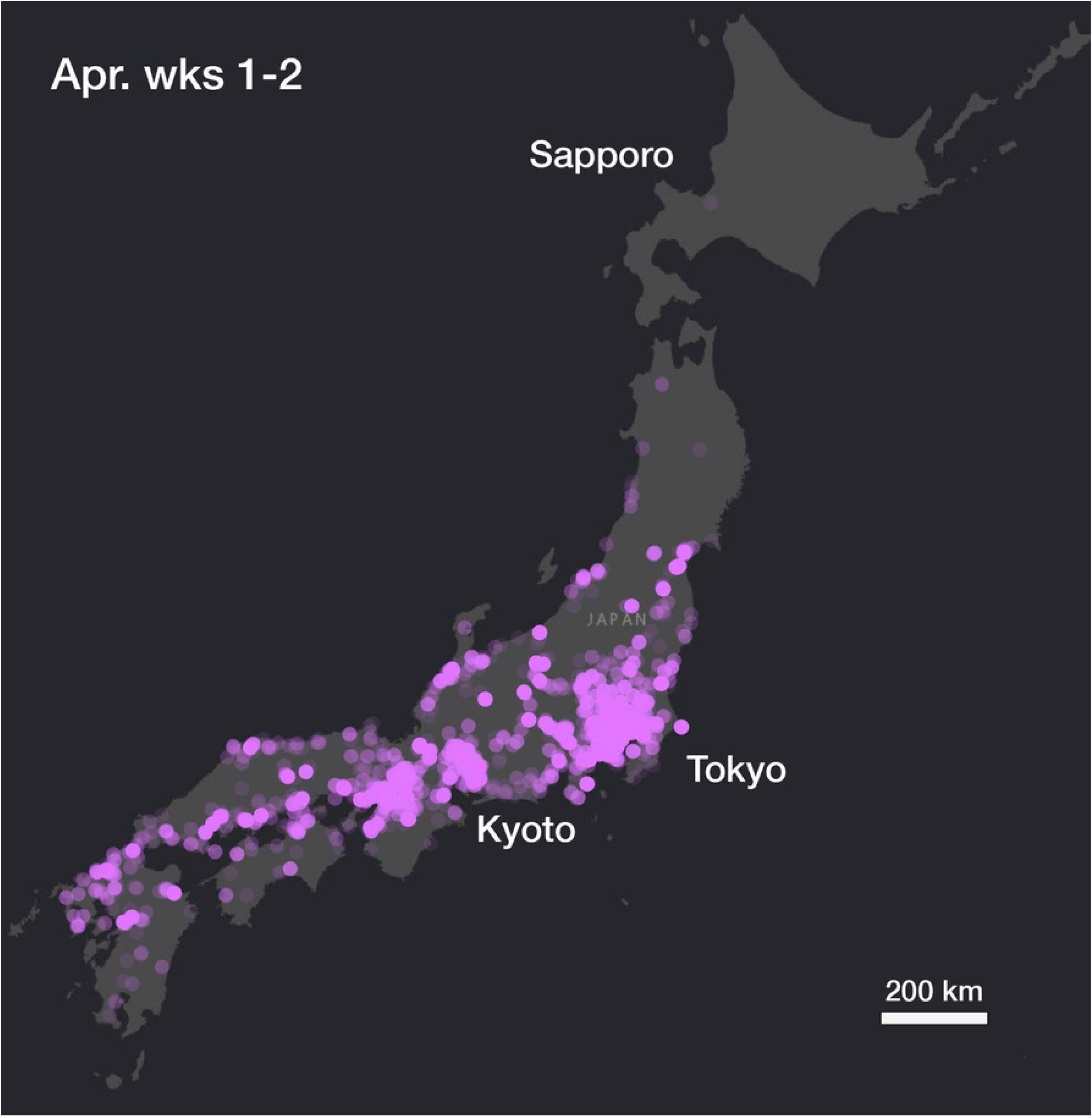
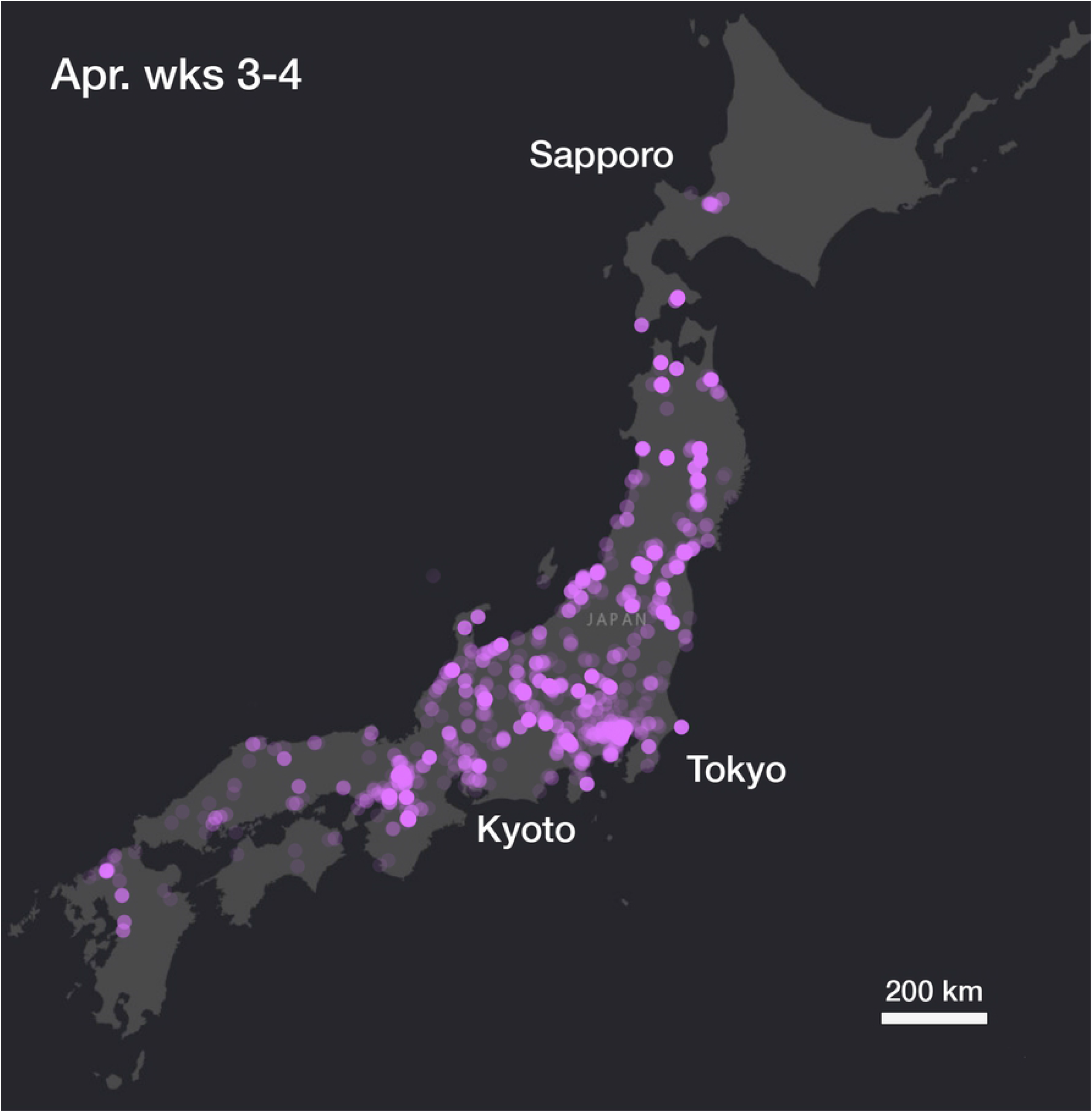
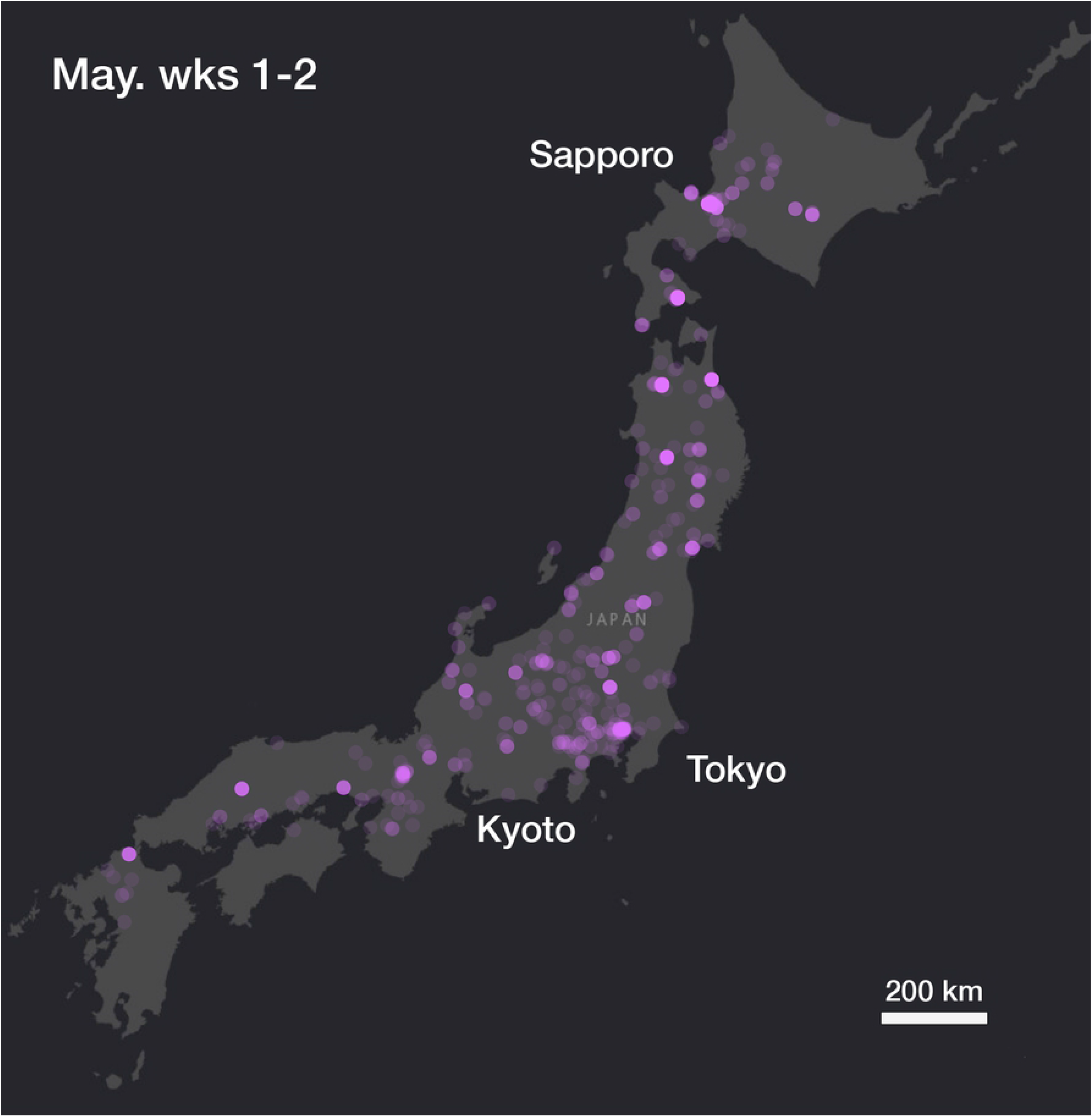
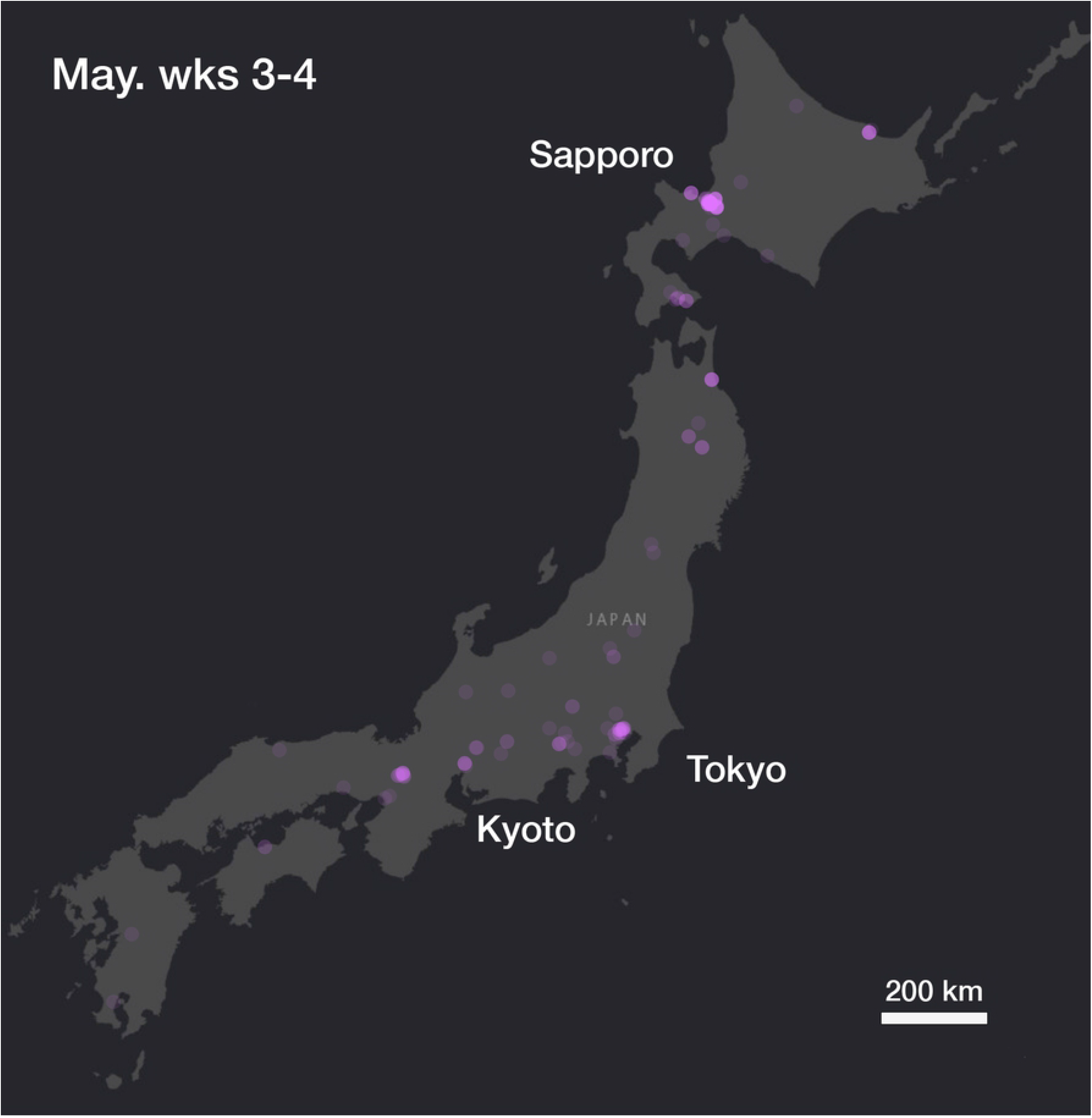
Cherry blossom photo hotspots in Japan from Flickr data 2008 – 2018. Each frame corresponds to a 2-week calendar period. **(A-C)** Cherry blossom photos appear in the warmer southern region of Japan, a higher concentration of photos shows in the urban centres of Tokyo and Kyoto on the main island of Honshu. **(D-F)** Cherry blossom photos intensify and begin to stretch northwards across the main island of Honshu. **(G-I)** Cherry blossom hotspots move north, appearing in the city of Sapporo on the northern island of Hokkaido. The urban centres of Tokyo and Kyoto remain hotspots, whilst blossom photography intensifies in Hokkaido and the north of Honshu. Finally, cherry blossom photos are subdued nation-wide moving upwards in a retreating wave from south to north. An animated/video sequence of the documented flower event is available as S1.

### 3.2 Tokyo image analysis

Our Flickr search for “cherry blossom” (section 2.1) returned 28,875 photos, geo-located within the district boundaries we used to identify the Tokyo region (Table 1, column B). The text tags, and their relative frequencies, returned by the machine vision API for this data set are reported (Fig 5). Out of the images returned from the text tag filter, 21,908 were subsequently tagged “cherry blossom” by the computer vision API (section 2.2), but some were eliminated due to them being tagged also “autumn” or “maple tree”, resulting in a set of 21,633 images (Table 1, column C). From the resulting images, the random sample selected for human checking is listed (Table 1, column D) and of these, the proportion confirmed to be cherry blossoms is listed also (Table 1, column E). The estimated accuracy of our automated processing method using computer vision and tag analysis is provided for each month (Table 1, column F). The accuracy is used to calculate the total cherry blossom photos in February, March, and April (Table1, column G) by multiplying (Table 1, column C) x (Table 1, column F).

**Table 1.**
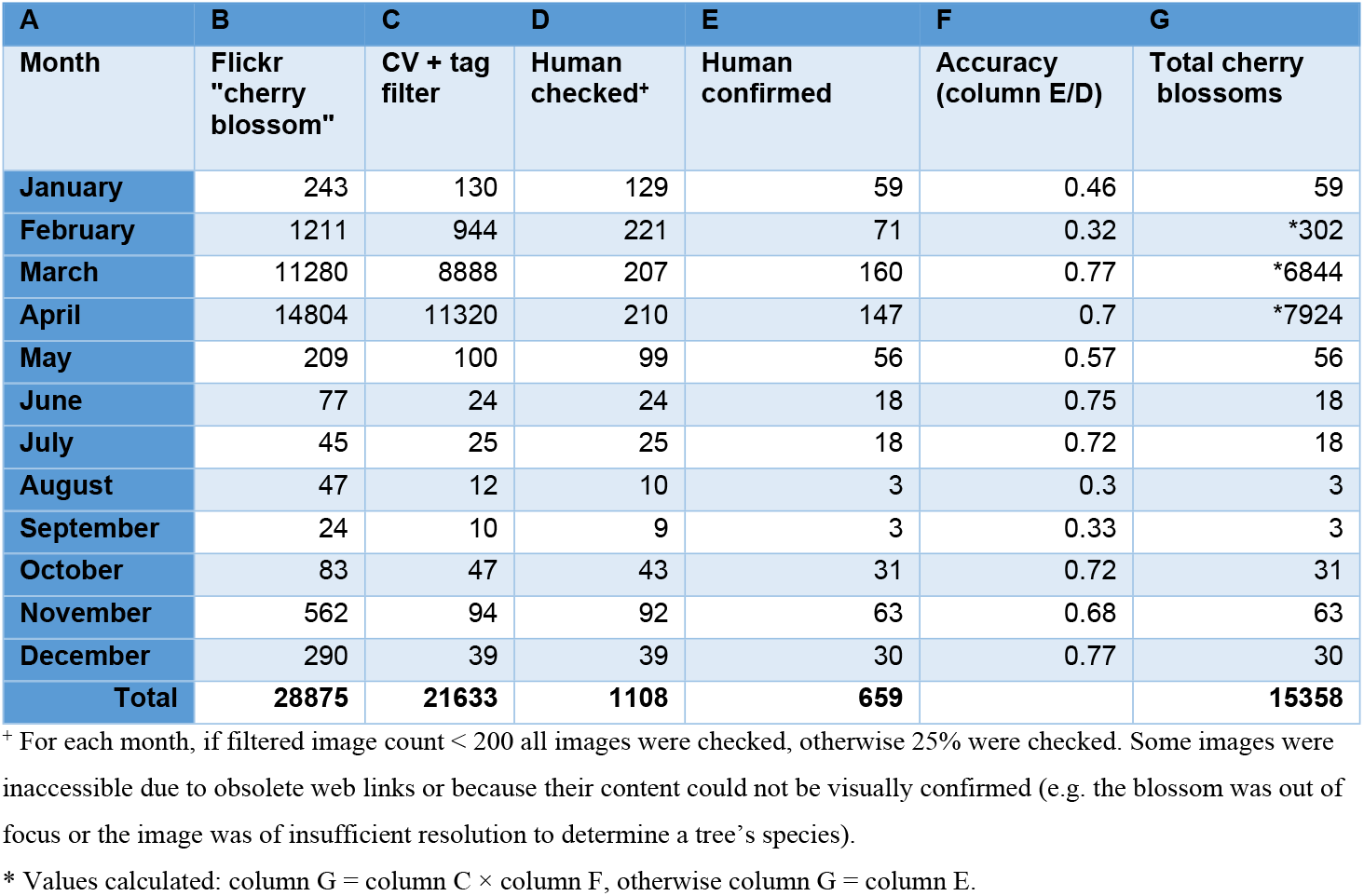
Image data counts in the Tokyo-filtered dataset at each stage of our process.

| A | B | C | D | E | F | G |
| --- | --- | --- | --- | --- | --- | --- |
| Month | Flickr "cherry blossom" | CV + tag filter | Human checked* | Human confirmed | Accuracy (column E/D) | Total cherry blossoms |
| January | 243 | 130 | 129 | 59 | 0.46 | 59 |
| February | 1211 | 944 | 221 | 71 | 0.32 | *302 |
| March | 11280 | 8888 | 207 | 160 | 0.77 | *6844 |
| April | 14804 | 11320 | 210 | 147 | 0.7 | *7924 |
| May | 209 | 100 | 99 | 56 | 0.57 | 56 |
| June | 77 | 24 | 24 | 18 | 0.75 | 18 |
| July | 45 | 25 | 25 | 18 | 0.72 | 18 |
| August | 47 | 12 | 10 | 3 | 0.3 | 3 |
| September | 24 | 10 | 9 | 3 | 0.33 | 3 |
| October | 83 | 47 | 43 | 31 | 0.72 | 31 |
| November | 562 | 94 | 92 | 63 | 0.68 | 63 |
| December | 290 | 39 | 39 | 30 | 0.77 | 30 |
| **Total** | **28875** | **21633** | **1108** | **659** |  | **15358** |

\* For each month, if filtered image count < 200 all images were checked, otherwise 25% were checked. Some images were inaccessible due to obsolete web links or because their content could not be visually confirmed (e.g. the blossom was out of focus or the image was of insufficient resolution to determine a tree's species).
\* Values calculated: column G = column C × column F, otherwise column G = column E.

**Fig 5.**
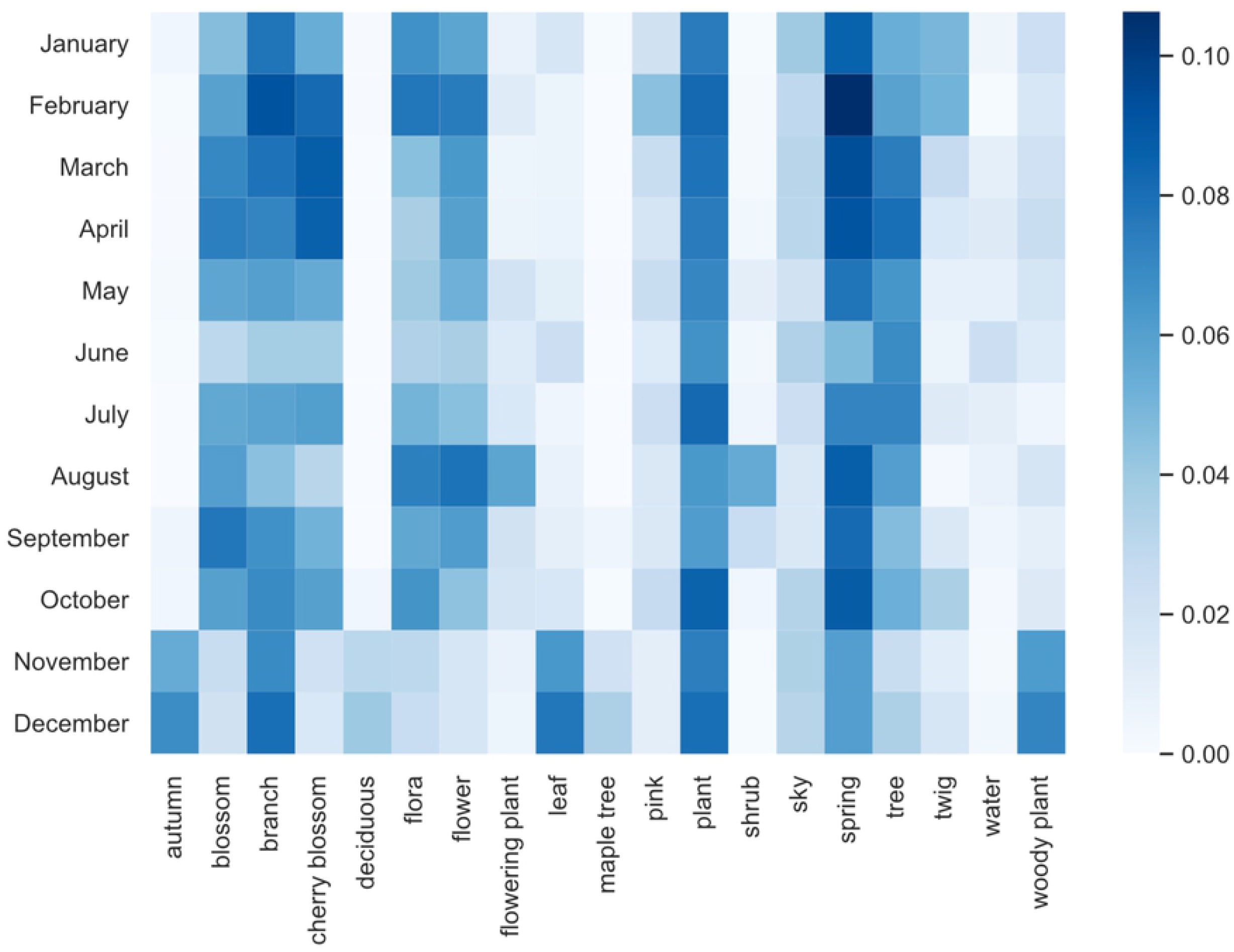
Normalised frequency of computer vision tags assigned to photos returned by Flickr search “cherry blossom”. Data for the Tokyo region over the period January 2008 to December 2018 inclusive. Images within tags are not mutually exclusive; an image may be labelled simultaneously by several tags. To generate this plot, the top-10 most frequent text tags returned for images in each month were collected into a set of 18 tags. These appear alphabetically along the x-axis. The square marked for each month against a tag is coloured according to the relative frequency of its prevalence in that month. Note that “autumn” is a frequent tag in November, and December. These photos were found to contain autumn leaves rather than blossoms.

### 3.3 Tokyo spring bloom estimation

The crests of the time series of our filtered and smoothed SNS cherry blossom photographic activity for the Tokyo region were calculated (Table 2). To validate these results, our SNS-estimated dates were compared against the published JNTO full bloom dates [40] and the resulting Root Mean Squared Error (RMSE) in our date was 3.21 days. The results of the comparison of our SNS-based estimate JNTO data for each year of the study period are plotted (Fig 6).

**Table 2.** Tokyo dates of Flickr SNS geotagged photo count peaks and JNTO-reported cherry full bloom dates. Peaks are computed on data smoothed using a 7-day triangular rolling average. JNTO data from [40].

| Year | Tokyo<br>SNS-estimated<br>full bloom | Tokyo<br>JNTO-reported<br>full bloom | Error (days) |
| --- | --- | --- | --- |
| 2018 | 26-Mar | 24-Mar | 2 |
| 2017 | 2-Apr | 2-Apr | 0 |
| 2016 | 3-Apr | 31-Mar | 3 |
| 2015 | 30-Mar | 29-Mar | 1 |
| 2014 | 31-Mar | 30-Mar | 1 |
| 2013 | 23-Mar | 22-Mar | 1 |
| 2012 | 8-Apr | 6-Apr | 2 |
| 2011 | 10-Apr | 6-Apr | 4 |
| 2010 | 9-Apr | 1-Apr | 8 |
| 2009 | 5-Apr | 2-Apr | 3 |
| 2008 | 29-Mar | 27-Mar | 2 |

**Fig 6.**
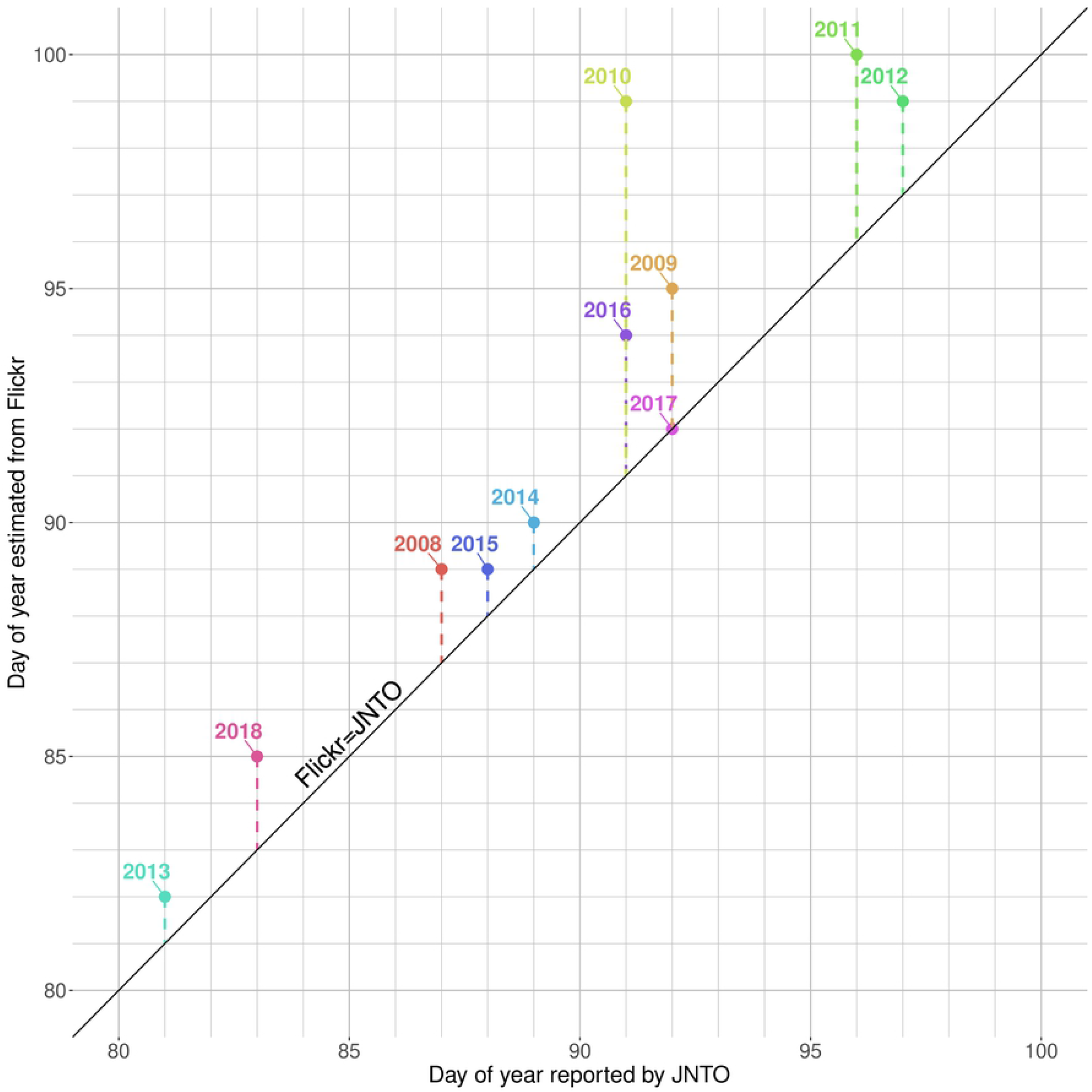
SNS-estimates for bloom date from Flickr data within the Tokyo region. The data (Table 2) is plotted as day number from 1^st^ January of each year during the study period 2008-2018. The black diagonal line represents the front along which SNS site Flickr and JNTO estimates are equal. The dashed lines extending from this diagonal represent the difference in number of days between our SNS-estimates and the JNTO-reported dates. Note that all SNS-estimates fall above or on the diagonal line, meaning that SNS activity lags or coincides with JNTO dates for reasons discussed below.

### 3.4 Tokyo autumn bloom identification

The temporal distribution of cherry blossom search results in Tokyo was amalgamated across all years in the study period by month and the monthly total calculated (Fig 7A). This data’s annual pattern has its main peak early in the year corresponding roughly to the Northern spring, as did the Japan-wide image set (*cf* Fig 1). In both the Japanese national and subset Tokyo-restricted datasets, a secondary peak of SNS images of cherry blossoms is apparent from October to December (Figs 7A & 7B).

**Fig 7.**
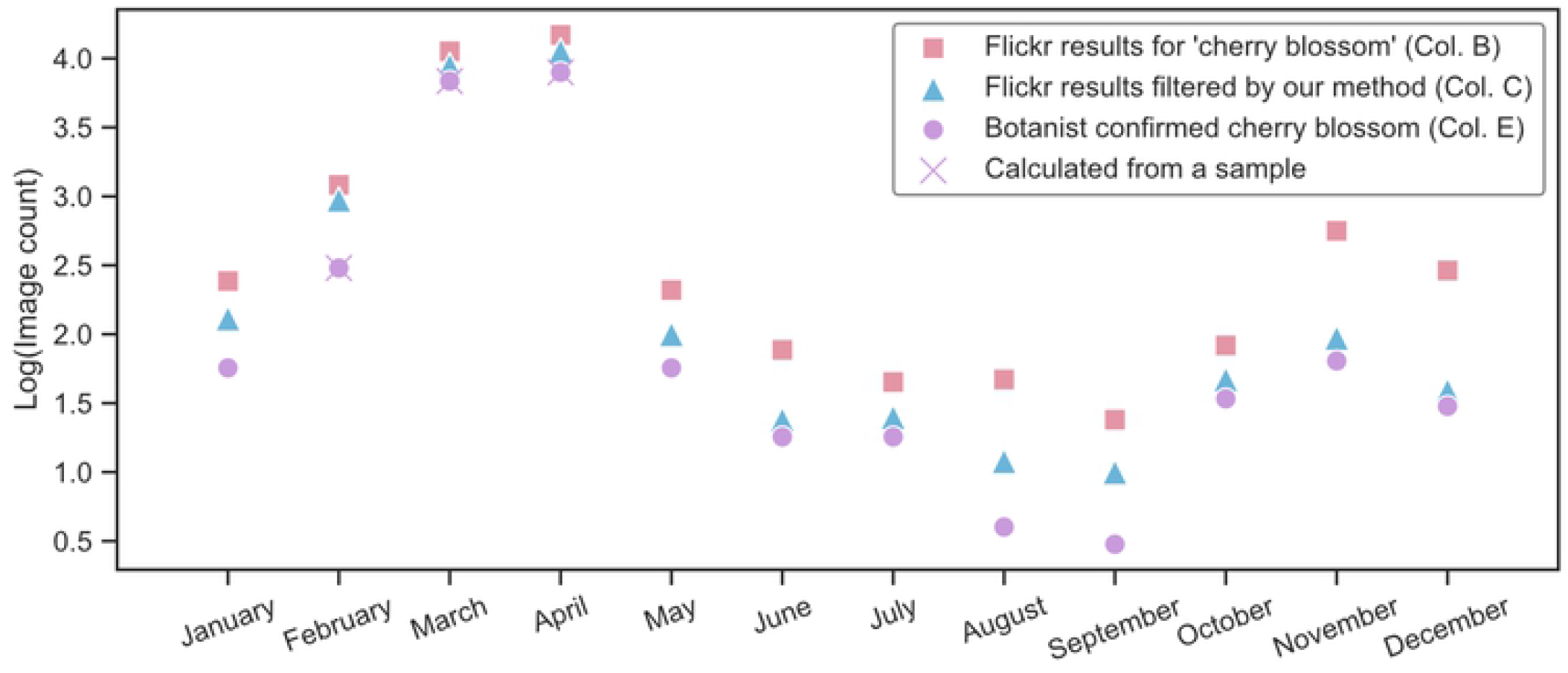
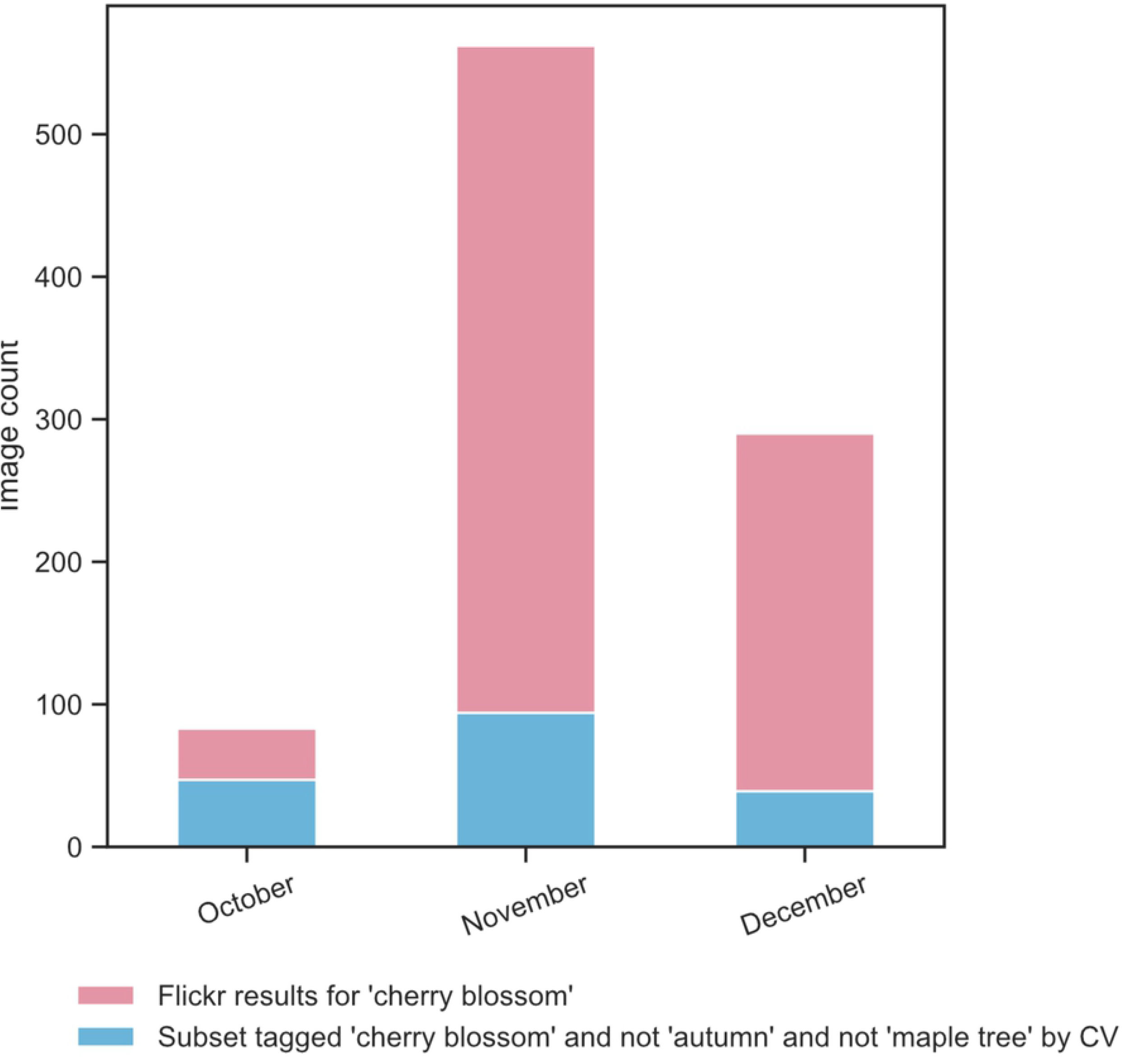
(A) Monthly total image count distribution of results from Flickr search “cherry blossom”. Data for Tokyo region 2008 – 2018 inclusive. Note y-axis uses a log-scale. Legend shows the data’s relationship to corresponding columns in Table 1. **(B)** Total count (pink + blue bars) of Flickr search results tagged “cherry blossom”. Data for Tokyo, for months October to December in the years 2008 – 2018 inclusive. A subset of these photos (blue bars) was also tagged “cherry blossom” and not “autumn” and not “maple tree”, by the computer vision API (See Table 1, column B).

The evidence for the November-centred flowering period was unexpected. Hence, to ensure that the photographs were not simply misclassified in this region due to an error in our method, we manually verified the content of the photos for months June to December for all years in our dataset. The monthly count of images confirmed to be cherry blossoms are plotted in Fig 7A (violet circles), demonstrating the veracity of the late flowering activity.

### 3.5 Kyoto spring bloom estimation

Our search for geotagged photos tagged “cherry blossoms” in the geographic boundary of Kyoto (gadm.org) during the study period January 2008 to December 2018 returned 14,516 results. Of these, 10,425 were tagged “cherry blossom”, and not “autumn”, and not “maple trees” by the computer vision API. The crests of the time series of our filtered and smoothed SNS cherry blossom photographic activity for the Kyoto region were calculated for each year of the study period (Table 3). To validate these results, our SNS-estimated dates were compared against the published JNTO full bloom dates [41], and the resulting Root Mean Squared Error (RMSE) in our date was 3.32 days.

**Table 3.** Kyoto dates of Flickr SNS geotagged photo count peaks and JNTO-reported cherry full bloom dates. Peaks are computed on data smoothed using a 7-day triangular rolling average. JNTO data from [41].

| Year | Kyoto<br>SNS-estimated<br>full bloom | Kyoto<br>JNTO-reported<br>full bloom | Error (days) |
| --- | --- | --- | --- |
| 2018 | 29-Mar | 28-Mar | 1 |
| 2017 | 7-Apr | 7-Apr | 0 |
| 2016 | 9-Apr | 2-Apr | 7 |
| 2015 | 4-Apr | 1-Apr | 3 |
| 2014 | 7-Apr | 2-Apr | 5 |
| 2013 | 31-Mar | 30-Mar | 1 |
| **2012** | 10-Apr | 9-Apr | 1 |
| **2011** | 8-Apr | 7-Apr | 1 |
| **2010** | 5-Apr | 1-Apr | 4 |
| **2009** | 8-Apr | 5-Apr | 3 |
| **2008** | 4-Apr | 1-Apr | 3 |

## 4 Discussion

In the current study we formally assess SNS images within Japan to understand if blooming patterns of cherry blossom trees can be reliably estimated from this data. Fig 4 evidences a shift in cherry blossoms nationwide from southern to northern Japan over 18 weeks, demonstrating a successful extraction of a national scale spatiotemporal phenological event from SNS data. We show that this event can be mapped with our technique to at least 2-week resolution, allowing a level of precise mapping with big data and no costly formal survey. Such precise nationwide data sampling of an important biological event was unimaginable until recent times. With growing concerns about the effects of climate change on plant phenology, and on changes to the spatiotemporal distributions of flowering plants and their pollinators [42, 43], incidental citizen science of the type we apply here is a powerful new tool to map the climate’s effect on our environment. The data also likely has benefits for tourism management and business activity planning.

Tokyo and Kyoto are suitable city-sized study sites for the application of SNS data to ecological phenomena, whereas other study sites, and other high-profile ecological phenomena besides cherry blooms, might be more difficult subjects for study. Firstly, their status as popular tourist destinations (Tokyo for instance is the eighth most-visited city in the world [44]) ensures that many geotagged photos of cherry blossoms are shared by visitors on popular SNS. Additionally, the Japanese National Tourism Office (JNTO) publishes the annual cherry blossom full bloom dates for the cities [40, 41], against which it was possible to validate our method. Finally, the cultural value Japan’s own citizens place on cherry blossoms, and on photography as a means to share cultural experiences, has long been documented [45]. Together, these characteristics assist in enlisting local residents, including Tokyo’s 9.27M people and Kyoto’s relatively small local population of 1.47 M (UN Data, 2015), as incidental citizen scientists. However, our data collection method also showed it is possible to capture the temporal signature of the cherry blossom event even in regions away from major cities (Fig. 4 and S1 Movie).

Applying our methodology, the SNS data can be successfully used to efficiently regenerate the dates of the flowering events to the JNTO-reported dates over a decade, in both Tokyo, and Kyoto. We note that our estimate each year for both regions consistently lagged or matched, but never foreshadowed, the date reported by JNTO (Tables 2 & 3). There may be a number of reasons for this. For instance, the SNS data peak may be generated as part of a self-actualising prophecy in which it follows the JNTO-published date. For this to be the case, for some reason a majority of visitors to the cherry blossoms would choose to visit only after published full bloom dates, biasing the SNS data in the way we observed. An ethnographic survey might elucidate the relevance of this effect on our project, but that is well beyond our present scope. However, we comment below on the extent to which our method stands independently of the JNTO-reported dates.

An alternative reason for the lag in SNS data with respect to the published dates might derive from some characteristic of the timestamping of images uploaded. Perhaps this might be biasing the SNS data somehow. For instance, perhaps international visitors to the site were more likely to have their phones set to a time zone preceding Tokyo’s rather than after it, ensuring that the images uploaded were frequently offset as we observed. We find this argument difficult to justify given the auto-update of mobile phone time zones connecting to the cellular network. It is also possible that photographic peaks are skewed by the weather – something that has been discussed elsewhere in the literature [10]. The first sunny day following the advertised bloom date might be the favoured time for visiting and photographing the cherry blossoms. Similarly, weekends and public holidays might be expected to impact photography patterns. As noted above, we smoothed data using a 7-day window to reduce the impact of weekly cycles (Fig 3). The only official Japanese public holiday in the vicinity of Hanami festivities is the Spring Equinox. This usually occurs 20^th^ or 21^st^ March each year and is therefore outside the range of both our predictions and the dates for peak bloom reported by JNTO. In addition, the holiday is not culturally associated with hanami.

Surprisingly, our analysis of the Flickr photos in Tokyo revealed the sensitivity of our technique by detecting evidence of a subtle, and initially unexpected (to the researchers at least), bloom in autumn (Figs 1, 3, 7). Although small, this peak’s authenticity was confirmed by manual inspection. The size of the peak could be indicative of several factors. Perhaps there were few photographs of the autumn bloom because there were few trees in bloom. Perhaps there were few photographs because few people knew about the bloom and/or few visited to photograph them. Autumn is not traditional for hanami and so, even if people did know through media and social networks about the bloom, they would have been unlikely to have made plans to visit and photograph the trees. This is especially true for international guests who must plan their travel well in advance and would probably target the main season. In short, the evidence of the late autumn bloom is likely to be doubly incidental in the sense that the “citizen scientists” were incidentally contributing ecological data, but also, they were unlikely to have deliberately set out to witness this phenomenon.

The autumn bloom has, it turns out, gained mainstream media attention: “for the first time in memory” [46, 47], “premature” [48], and “unexpected” [49]. Our data shows the phenomenon to have been evident in Tokyo for many years. This is certainly interesting in and of itself, but also, it is evidence that our method is not simply a demonstration of the self-fulfilling prophecy generated by the JNTO-published dates –this secondary peak is not officially noted by JNTO yet our tools indicate its presence within the data.

One possibility is that the extra bloom is due to the presence of cherries of species *P. Subhirtella*, known to flower in colder weather [50]. However, the cited news outlets have proposed the event to be a possible outcome of climate change. It is well beyond the scope of our study to determine the relative merits of these two proposed causes. However, the merit of our SNS image analysis methodology is that it can detect such fine-grained shifts in flowering plant behaviour and spark further investigations.

Our results show that in the photo set we collected using a Flickr search for “cherry blossoms”, photos in January and February were falsely interpreted by our method as cherry blossoms (only 44% and 32% were correct respectively). This may be attributed to photos of plums, which precede cherries as they bloom in colder weather [51], often whilst snow is still present. So, plum blossoms are expected to be concentrated in this time of the year and may be confused with cherry blossoms. Accuracy is also low in July and August. In these months, cherries are not in bloom, and very few photos exist for these months during the period of our study. It is worth noting that the distance between the phenological front of plum and cherry bloom times varies across the length of Japan [52]. Therefore, the classification error may be expected to change depending on the region under study. For instance, plums and cherries may flower simultaneously in Hokkaido in the North, while in the Southern island of Kyushu the difference in blooming dates may be wide.

Recently, computer vision algorithms have moved from focussing on broad object categories (e.g. plant, car, etc.) to fine-grained visual recognition competitions (FGVCs) where the objective is to differentiate between sub-categories within broad classifications [53]. When API providers like Google participate in these activities [54] the results can improve workflows like ours, since APIs for species classification may identify and distinguish between close species (such as cherry/plum blossoms) where visual distinctions between their morphology exist.

Our results show that the SNS data can give valuable insights into phenological activities. However, we must be careful when drawing conclusions about the relative densities of photographs in space and time; it has been noted in the literature that SNS geo-tagged photo content may be spatially and temporally biased, not only by the underlying physical subject of the content, but also by human photographic activity [10]. In a part of the current work, we localised our temporal analysis of cherry blossoms to the cities of Tokyo and Kyoto. This is one way to minimise the variation in human population density across a region of study (and the ensuing variation in photo density) as well as to minimise the physical climate variation. The popularity of Tokyo and Kyoto as global tourist destinations, and their resident density, makes this feasible. In addition to the temporal analysis for the two cities specifically, we collected and analysed data geo-localised to Japan in its entirety. This demonstrates that within the limits of a data unnormalized for population density, it is still valuable to apply techniques like those studied here across nationwide data sets (e.g. see [10]).

It is technically more challenging, but potentially still worthwhile, to use the Japanese character in the Flickr search API since it is a Unicode, not ASCII character. The Japanese character for cherry blossom 桜 (Sakura) was therefore tested on Flickr and we found it returned a smaller result set than the English “cherry blossom”. Nevertheless, to ensure we did not overstep our linguistic capabilities, we opted for the English search keyword in this demonstration study since it does not compromise our intent. For a complete survey of a phenomenon in an ecological context we suggest it would be worthwhile to include data text-tagged in the relevant local language, including common names and colloquial terminology. The research community in this area has discussed the use of scientific versus common names to search for species also (e.g. [10, 13]).

While analysing photo content, we excluded photos (Table 1, labelled ‘+’) in cases where it was difficult to visually confirm whether they were of cherry blossoms. In the future, this decision can be enhanced by exploiting location information and cross-checking against other sources (e.g. Google street view in the same location, or photos from other social networks) especially in urban areas where there is high photographic activity.

## 5 Conclusions

The Japanese cherry bloom is the quintessential public ecological phenomenon. Its status in Japan as a cultural event ensures local attention. A global fascination with Japanese culture also ensures tourists flock to see the blossoms and experience the Hanami cherry blossom viewing festivities. Our study indicates that with our processing and filtration method, social media data sources are capable of capturing the spatiotemporal sweep of cherry blossoms across the country at a fine scale. Our method therefore promises to enable new insights on a variety of similarly photogenic phenomena. In the current study we also use this powerful new technique to trace cherry peak bloom over a decade in two Japanese cities, Tokyo and Kyoto. Our results were consistently accurate to within a few days of the official Japanese Tourism organisation’s reported full bloom dates for these cities. Finally, we showed that social network data has the power to reveal seemingly subtle anomalies in phenological phenomena such as out-of-season flowering times.

Overall, our findings suggest that our methods have the potential to help us understand changing ecological phenomena in a world that is increasingly putting its data online for all to see. Our techniques can provide one means to fill gaps in published data that will help us understand the impact of the changing climate on our ecosystems.

## SUPPLEMENTARY MATERIALS - CAPTIONS

**S1 Movie. An animation of cherry blossom photographic hotspots, Japan 2008-2018.**

An animation in which each frame corresponds to a 2-week calendar period of cherry blossom photography activity across Japan during the decade 2008-2018. Each year, at first, cherry blossom photos appear in the warmer southern region of Japan. Higher concentration of photos shows in the urban centres of Tokyo and Kyoto on the main island of Honshu. Next, cherry blossom photos intensify and begin to stretch northwards across the main island of Honshu. Then, cherry blossom hotspots move north, appearing in the city of Sapporo on the northern island of Hokkaido. The urban centres of Tokyo and Kyoto remain hotspots, whilst blossom photography intensifies in Hokkaido and the north of Honshu. Finally, cherry blossom photos are subdued nation-wide moving upwards in a retreating wave from south to north.

**S1 Data. Flickr image ids and associated metadata**

Contains data obtained from Flickr, filtered via computer vision-generated text tags.

Fields are:

-id:Photo Id in Flickr
-url:URL to the photo on Flickr
-visual_tags:Text tags generated for the photo via computer vision
-dateTaken:Date the photo was taken
-lat:Latitude of the photo location
-lng:Longitude of the photo location

**S2 Data. Human-visual cherry blossom image review data**

Contains photos manually reviewed by a botanist to confirm whether they actually are cherry blossoms

Fields are:

-id:Photo Id in Flickr
-Q1:Does the photo include a plant/tree?
-Q2:Does the plant/tree have flowers?
-Q3:Are flowers identifiable as cherry blossoms?

